# Absence of Genome Reduction In Diverse, Facultative Endohyphal Bacteria

**DOI:** 10.1101/045708

**Authors:** David A. Baltrus, Kevin Dougherty, Kayla R. Arendt, Marcel Huntemann, Alicia Clum, Manoj Pillay, Krishnaveni Palaniappan, Neha Varghese, Natalia Mikhailova, Dimitrios Stamatis, T. B. K. Reddy, Chew Yee Ngan, Chris Daum, Nicole Shapiro, Victor Markowitz, Natalia Ivanova, Nikos Kyrpides, Tanja Woyke, A. Elizabeth Arnold

## Abstract

Fungi interact closely with bacteria both on the surfaces of hyphae, and within their living tissues (i.e., endohyphal bacteria, EHB). These EHB can be obligate or facultative symbionts, and can mediate a diverse phenotypic traits in their hosts. Although EHB have been observed in many major lineages of fungi, it remains unclear how widespread and general these associations are, and whether there are unifying ecological and genomic features found across all EHB strains. We cultured 11 bacterial strains after they emerged from the hyphae of diverse Ascomycota that were isolated as foliar endophytes of cupressaceous trees, and generated nearly complete genome sequences for all. Unlike the genomes of largely obligate EHB, genomes of these facultative EHB resemble those of closely related strains isolated from environmental sources. Although all analyzed genomes encode structures that can be used to interact with eukaryotic hosts, we find no known pathways that facilitate intimate EHB-fungal interactions in all strains. We isolated two strains with nearly identical genomes from different classes of fungi, consistent with previous suggestions of horizontal transfer of EHB across endophytic hosts. Because bacteria are differentially present during the fungal life cycle, these genomes could shed light on the mechanisms of plant growth promotion by fungal endophytes during the symbiotic phase as well as degradation of plant material during saprotrophic and reproductive phases. Given the capacity of EHB to influence fungal phenotypes, these findings illuminate a new dimension of fungal biodiversity.

## Introduction

All eukaryotes have evolved in the presence of bacteria, with diverse bacteria adopting an endosymbiotic and intracellular habitat across the eukaryotic tree of life (Sachs *et al*., 2011; Schulz & Horn, 2015). Much like the diverse Metazoa that host rich bacterial microbiomes, fungi interact closely with bacteria both on the surfaces of hyphae, and within their living tissue (i.e., endofungal or endohyphal bacteria, EHB) (Arendt *et al*., 2016; Hoffman & Arnold, 2010; Naito *et al*., 2015; Partida-Martinez & Hertweck, 2005; Torres-Cortés *et al*., 2015). These EHB can be either obligate or facultative symbionts and can mediate diverse phenotypic traits in their hosts. For instance, EHB inhabiting some rhizosphere fungi can influence virulence of phytopathogens or the capacity of mycorrhizal fungi to establish symbiotic associations (Partida-Martinez & Hertweck, 2005; Salvioli *et al*., 2015). In turn, EHB inhabiting foliar fungal endophytes (fungi that occur in living leaves without causing disease; class 3 endophytes, sensu Rodriguez *et al*., 2009) can increase the production of plant growth-promoting hormones (Hoffman *et al*., 2013) and alter the capacity of their hosts to degrade plant tissues (Arendt, 2015). Although EHB have been observed in many of the major lineages of plant-associated fungi (including diverse Mucoromycotina, Glomeromycota, Basidiomycota, and Ascomycota), it remains unclear how widespread and general these associations are across important fungal species, and whether there are unifying ecological and genomic trends found among all EHB strains. To better understand genomic characteristics of facultative EHB associated with fungal endophytes, we isolated 11 bacterial strains after they emerged from the hyphae of diverse Ascomycota that were isolated as endophytes of cupressaceous plants, and generated nearly complete genome sequences for all.

Multiple EHB associated with other fungal taxa have already been sequenced and analyzed in genome-level studies (e.g., Lackner *et al*., 2011; Naito *et al*., 2015; Torres-Cortés *et al*., 2015), providing a framework for determining whether previously identified genomic trends hold across all endohyphal bacterial symbionts. *Burkholderia rhizoxinica* inhabits hyphae of the plant pathogen *Rhizopus microsporus,* where this bacterium produces a toxin required for fungal pathogenicity on rice (Partida-Martinez & Hertweck, 2005). Notably, this symbiosis is established and maintained by the presence of a type III secretion system, although precise effector genes are not known at this time (Lackner *et al*., 2010). Because *B. rhizoxinica* can be vertically transmitted through fungal spores, the close association of fungal host and bacterial symbiont across generations enables and may even select for genome reduction as seen in other obligate intracellular symbionts (Delaye, 2015). Similarly, the genomes of mollicutes-related endobacteria (MREs), which are also obligate symbionts, are relatively small compared to those of environmental bacteria (Naito *et al*., 2015; Torres-Cortés *et al*., 2015). However, genomes of Mollicutes are generally small, such that size reductions may have occurred before evolution of the endofungal lifestyle (Barre, 2004). Alternatively, because the trend for genome reduction in obligate symbionts is quite strong, genome size relative to outgroups may be used as a proxy for mode of transmission.

In contrast to the genomes of previously studied, largely obligate EHB, here we find that the sizes of genomes from facultative EHB resemble those of closely related strains isolated from environmental sources. While these EHB strains all possess structures that are capable of interacting with eukaryotic hosts, we do not find evidence for a conserved pathway that mediates EHB-fungal interactions across all strains. We consider these genome data to be informative regarding little-known aspects of the transmission and population dynamics of EHB. Because these bacteria can influence plant growth promotion by fungal endophytes during the symbiotic phase (Hoffman & Arnold, 2010) as well as degradation of plant material during saprotrophic and reproductive phases (Arendt, 2015), these genomes could enable engineering of symbiotic associations to enhance growth and processing of plant material.

## Materials and Methods

### Isolation of Bacterial Strains and Genomic DNA

To trigger emergence of bacterial strains from their fungal hosts, mycelia were grown from mycelial plugs on 2% malt extract agar at 36°C (Hoffman & Arnold, 2010; Hoffman *et al*., 2013; Arendt *et al*., 2016). After 72 h, bacteria generally emerged from apparently axenic mycelium. Endohyphal status of bacteria was confirmed prior to emergence following Hoffman *et* al.,(2010) and Arendt *et* al.,(2016). Emergent bacteria were streaked to single colonies on LB media without antibiotic supplements. Individual colonies were grown in liquid LB media and frozen in 40% glycerol. Bacterial strains and genomic DNA were verified through PCR and Sanger sequencing of the 16s rRNA locus using universal primers 27F and 1492R (see Hoffman *et al*., 2010). Before isolating genomic DNA, bacterial strains were streaked from frozen stocks, at which point a single colony was inoculated into 5mL LB media and grown at 27^°^C overnight. Genomic DNA from this 5mL culture was isolated using a Wizard genomic DNA isolation kit following the manufacturer’s instructions (Promega, Madison WI).

### Genome Sequencing and Assembly

Draft and complete genomes were generated at the DOE Joint Genome Institute (JGI) using the Pacific Biosciences (PacBio) sequencing technology (Eid *et al*., 2009). A Pacbio SMRTbell™ library was constructed and sequenced on the PacBio RS platform.Characteristics of each sequencing run and assembly can be found in Table 1. All general aspects of library construction and sequencing performed at the JGI can be found at http://www.jgi.doe.gov. Raw reads were assembled using HGAP (version: 2.2.0.p1) (Chin *et al*., 2013).

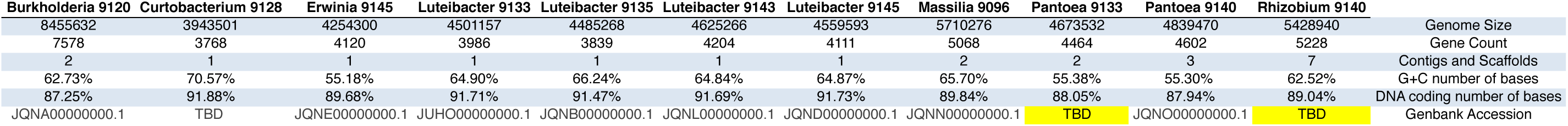

### Genome Annotation

Genomes were annotated using the JGI microbial annotation pipeline (Huntemann *et al*., 2015), followed by a round of manual curation using GenePRIMP (Pati *et al*., 2010) for finished genomes and draft genomes in fewer than 10 scaffolds. Predicted CDSs were translated and used to search the National Center for Biotechnology Information (NCBI) nonredundant database, UniProt, TIGRFam, Pfam, KEGG, COG, and InterPro databases. The tRNAScanSE tool (Lowe & Eddy, 1997) was used to find tRNA genes, whereas ribosomal RNA genes were found by searches against models of ribosomal RNA genes built from SILVA (Pruesse *et al*., 2007). Other non-coding RNAs such as the RNA components of the protein secretion complex and the RNase P were identified by searching the genome for the corresponding Rfam profiles using INFERNAL (http://infernal.janelia.org). Additional gene prediction analysis and manual functional annotation was performed within the Integrated Microbial Genomes (IMG) platform (http://img.jgi.doe.gov) developed by JGI, Walnut Creek, CA, USA (Markowitz *et al*., 2009). All additional genomic analyses, including pathway presence and absence, were carried out using the IMG platform.

### Phylogenetic and Comparative Genomic Analyses

Whole genome files for all strains listed in Figure 1 are publicly available at GenBank (see Table 1), with sequencing and assembly reports listed on Figshare at the following link: https://dx.doi.org/10.6084/m9.figshare.2759821.v1

**Figure 1.**
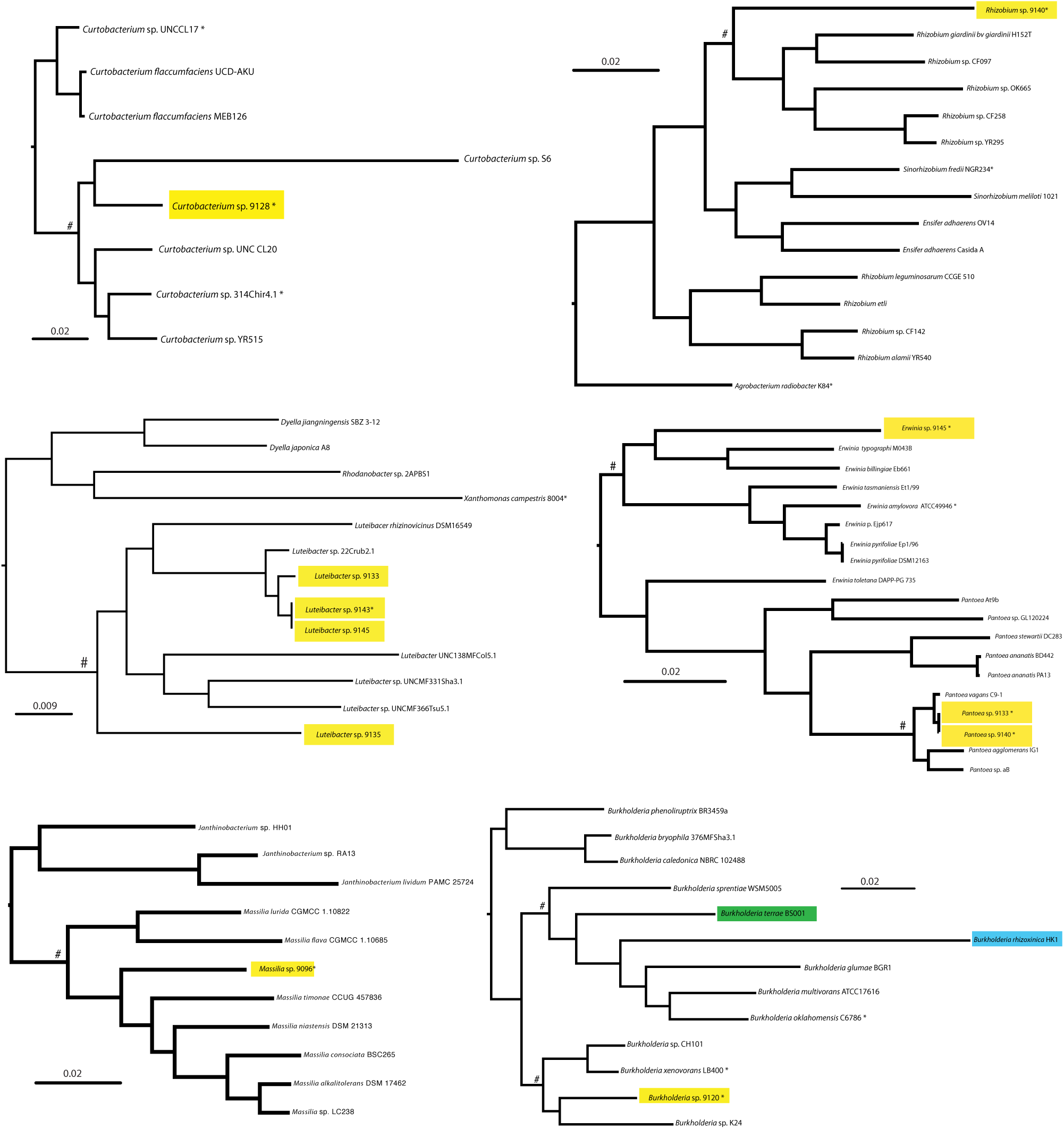
Phylogenetic Profiling Demonstrates Convergent Evolution of Facultative Endohyphal Bacteria. Maximum likelihood phylogenies for each of the focal EHB as well as closely related bacteria were built from whole genome sequences using RealPhy. Focal EHB strains are highlighted in yellow. Genomes for additional fungal-associated bacteria are highlighted in green (endohyphal) or blue (extracellular). Strains used as references for alignment in RealPhy are indicated by *, and the root node for strains used in genome size calculations (Figure 2) are denoted by “#”.

Phylogenies were constructed using the RealPhy online server (Bertels *et al*., 2014). Genbank accession numbers, or JGI accession identification numbers when genomes were not found on Genbank, can be found on Figshare at https://dx.doi.org/10.6084/m9.figshare.3124006.v1. Briefly, for each of phylogeny shown in Figure 1, GenBank files were uploaded to the server and maximum likelihood phylogenies were built from whole genome alignments to a single reference genome. Reference phylogenies were built to all strains denoted with “*” and then merged to produce the final phylogeny.

Geneious version 6.0.5 (Kearse *et al*., 2012) was used to compare whole genome alignments for *Erwinia* sp. 9140 and *Erwinia* sp. 9145, as well as *Luteibacter* sp. 9143 and *Luteibacter* sp. 9145. Briefly, sequences from these genomes were imported into Geneious and aligned using the Mauve option with default parameters. SNPs and indels were displayed as disagreements between these alignments, were visually inspected for proper alignment, and were counted by hand.

## Results and Discussion

### Convergent Evolution of Closely Related Endohyphal Bacteria

Because genome sequences typically provide a broader picture of evolutionary relationships among bacterial strains than phylogenies built from single loci, we inferred phylogenies based on whole-genome data generated for this study as well as closely related outgroups (Baltrus *et al*., 2014; Bertels *et al*., 2014). Our evaluation of multiple EHB strains from diverse Ascomycota provides insights into phylogenetic signals associated with the EHB lifestyle, an opportunity to explore shared genomic architecture relevant to the EHB lifestyle, and the opportunity to evaluate whether these EHB to have undergone convergent evolution.

For most of our focal EHB, the data suggest that the facultative endohyphal lifestyle has evolved multiple times amongst closely related bacteria. For instance, we found phylogenetically distinct strains of *Erwinia* and *Pantoea* within different classes of fungal hosts. The *Pantoea* strains are both close relatives of *Pantoea vagans,* a plant-associated epiphyte, but are also closely related to a known plant pathogen, *Pantoea agglomerans*. These relationships raise the intriguing possibility that some phytopathogens could potentially “hide out” inside of endophytic fungi, an hypothesis to be addressed in future experiments. In comparison, the *Erwinia* strain is placed outside of groups consisting of previously well-characterized genomes. Furthermore, our data demonstrate that the *Burkholderia* strain evaluated here is phylogenetically distinct from the two previously characterized *Burkholderia* and *Glomeribacter* strains and is significantly diverged from *Burkholderia terrae,* which forms a close relationship with fungi from soil (Nazir *et al*., 2012; van Elsas, 2014). Our *Rhizobium, Curtobacterium,* and *Massilia* isolates are the first from these clades to be found as EHB, although all are closely related to strains found associated with plants and throughout the environment. It remains a possibility that many different environmental bacteria can associate within fungal endophytes and therefore transiently be categorized as EHB, and in this case any convergent phylogenetic signals could represent sampling bias associated with our strains. For example, if signals of convergence were instead due to biases, we predict that we would find a diverse array of *Pantoea* and *Erwinia* species with more intense sampling of EHB from fungal populations. However, we note that our previous categorization based solely on 16s rRNA across a wider variety of fungal hosts also shows clustering of EHB strains into particular clades rather than the presence of diverse sequences from throughout the *Pantoea/Erwinia* phylogeny (Hoffman & Arnold, 2010).

Compared to the genomes of other facultative EHB, *Luteibacter* strains display an interesting phylogenetic pattern that suggests some level of host specificity. There are two distinct clades within the *Luteibacter* phylogeny: one that is mainly composed of rhizosphere isolates and one that is composed mainly of EHB. An additional EHB *Luteibacter* (strain 9135) appears as an outgroup to both clades. It is unclear whether this pattern differs from those found within other bacterial genera because of sampling biases, or whether the association of these *Luteibacter* strains with fungal hosts is truly distinct and specialized.

### Diverse Fungi Harbor Similar Symbionts

Most of the host fungi of these bacterial strains were isolated in from a small number of closely spaced trees in Duke Forest (Durham, North Carolina, USA) (Hoffman &Arnold, 2010). All of the focal EHB strains were isolated as they emerged from fungal cultures, and in some cases we isolated strains that were indistinguishable at the 16s rRNA level, yet occurred in phylogenetically divergent fungi. Whole genome sequencing can shed light on whether these EHB strains are members of the same clonal group or are just closely related isolates. We also note that it is possible that these strains represent colonization events within the laboratory environment, but the probability of such contamination is very low due to multiple measures employed to prevent and account for such incidents (see Arendt *et al*., 2016).

In the case of *Luteibacter* spp. 9143 and 9145, we found no verified SNPs that could distinguish their genomes from one another. Although 18 regions differ at a single nucleotide resolution between these two strains, all lie within homopolymer tracts and are therefore likely the product of sequencing errors. Some of these potential errors alter automatic annotation of the genomes and may account for many of the presumed differences in protein content between the strains. Because these two strains were isolated from highly divergent classes of Ascomycota (Dothideomycetes and Sordariomycetes, respectively), the lack of nucleotide diversity suggests horizontal transfer of these strains in nature. However, we also find that one 40,413bp region is present within *Luteibacter* sp. 9143 yet missing from the assembly of *Luteibacter* sp. 9145. This region encodes many phage-associated genes, and therefore likely encodes a prophage. It remains to be seen how the prophage affects the physiology of these strains.

In one additional case, we isolated similar EHB strains from diverse fungal hosts. However, we observe more diversity between *Pantoea* sp. 9140 and *Pantoea* sp. 9133 than between the *Luteibacter* strains mentioned above: we find 21 SNPs across conserved regions and alignable regions. Moreover, 10 of these SNPs appear to be true nucleotide polymorphisms because they are not associated with repetitive nucleotide tracts. *Pantoea* sp. 9140 contains additional sequences (11,970bp on one contig, 171,396bp on a separate contig) that do not appear to be present in *Pantoea* sp. 9133. Taken together, comparison of these strains further demonstrates that closely related bacteria can be found across divergent fungi, consistent with the lack of strict-sense cocladogenesis observed with natural hosts (see Hoffman & Arnold 2010; see also Arendt *et al*., 2016).

### Genomes of These Endohyphal Bacteria are Not Reduced

The genome sizes for many intracellular bacteria, including most known EHB, are drastically smaller than those of closely related free-living species (e.g., Ghignone *et al*., 2011; Lackner *et al*., 2011). Reductions in genome size are thought to be a product of reduced selection pressures on deleterious mutations due to repeated population bottlenecks, a deletion bias for bacterial genomes, and lack of selection to maintain physiological pathways made redundant because they are encoded by the host (Delaye, 2015). It is also possible that genomes may be directly streamlined by natural selection as a way to optimize metabolic efficiency (Grote *et al*., 2012). As such, a reduction in genome size compared to closely related bacteria speaks to ecological and evolutionary pressures experienced by intracellular bacteria and can therefore illuminate the bacterial lifestyle.

We compared the genome size of 11 EHB to closely related outgroup strains to test for reduction of genome size associated with the EHB lifestyle (Fig. 2). We also included the genome of *B. rhizoxinica* and used the same comparisons to demonstrate the signal for a known instance of genome reduction. In all but one case, genome sizes for our focal EHB fall either within or just below the range of genome sizes for these outgroups. The one strain that stands out, *Rhizobium* sp. 9140, possesses a genome 1Mb smaller than other closely related bacteria. We therefore see little evidence that these EHB have generally experienced widespread genome reduction.

**Figure 2.**
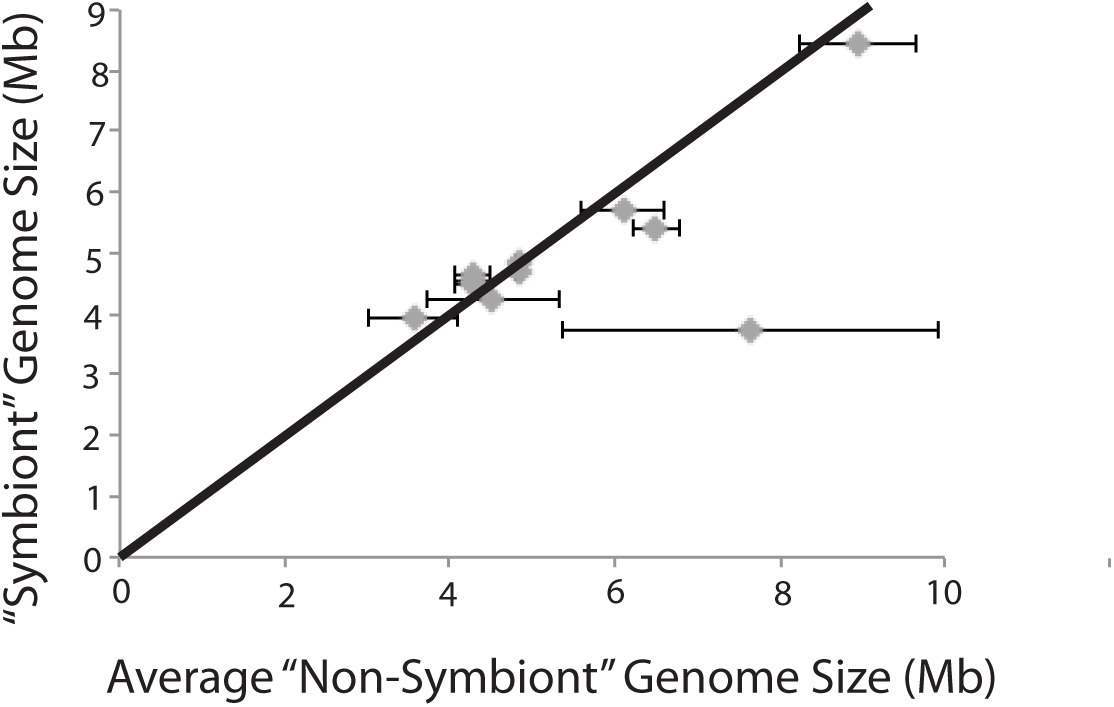
Absence of Genome Reduction in Facultative Endohyphal Bacteria. Whole genome sizes for each of the focal EHB strains are plotted on the Y axis, while the average genome sizes for a diverse suite of related bacteria (the root of all analyzed strains indicated by “#” in Figure 1) are plotted on the X axis. Error bars indicate 1 standard deviation for the “Non-symbiont” bacteria against which each “Symbiont” genome was plotted.

The absence of genome reduction within these strains is consistent with laboratory studies suggesting that they are gained and lost readily, that fungi are capable of major metabolic activity in the absence of the bacteria, that the bacteria can be isolated on standard laboratory media, and that they are transmitted horizontally (see Hoffman & Arnold 2010, Hoffman *et al*., 2013, Arendt *et al*., 2016). It is therefore possible that populations of these strains do not experience drastic population bottlenecks during transmission, and that diverse genomic architecture needed for survival outside of hosts has been maintained.

### Absence of Conserved Systems Known to Direct Intimate Interkingdom Interactions

In established systems of bacterial-fungal symbiosis, intimate interactions are usually carried out through the action of various bacterial secretion systems (Lackner *et al*., 2010). Indirect interactions are carried out in Gram negative and Gram positive bacteria by Type I, II, and V secretion systems, which secrete substrates outside of cells (Costa *et al*., 2015). Increasingly intimate interactions are largely carried out in Gram negative bacteria through the actions of type III, IV, and VI secretion systems, which are known to translocate substrates (effector proteins) directly into recipient cells (Costa *et al*., 2015). Both type II (for the secretion of chitinase) and type III secretion systems have been implicated in the establishment and maintenance of the *Burkholderia-Rhizopus* interaction (Lackner *et al*., 2010; 2011). Likewise, type III, IV, and VI secretion systems are important in interactions between bacteria and single-celled eukaryotes such as amoebae (Burstein, 2016; Matz *et al*., 2008; Van der Henst *et al*., 2015).

We queried all 11 complete genomes for evidence of secretion systems possibly involved in establishment of fungal symbiosis using JGI’s online annotation tools (Figure 3). General secretion pathways (types I and II) are likely found within all of these genomes, as expected based on their general presence across a majority of Gram negative bacteria isolated in culture (Costa *et al*., 2015; Delepelaire, 2004; Korotkov *et al*., 2012). Almost all strains except *Rhizobium* and *Curtobacterium* appear to encode basic type I systems. All 11 bacteria appear to encode both the Sec and Tat translocation systems, whereas only a subset of these have the genetic potential to create outer membrane proteins associated with type II secretion.

**Figure 3.**
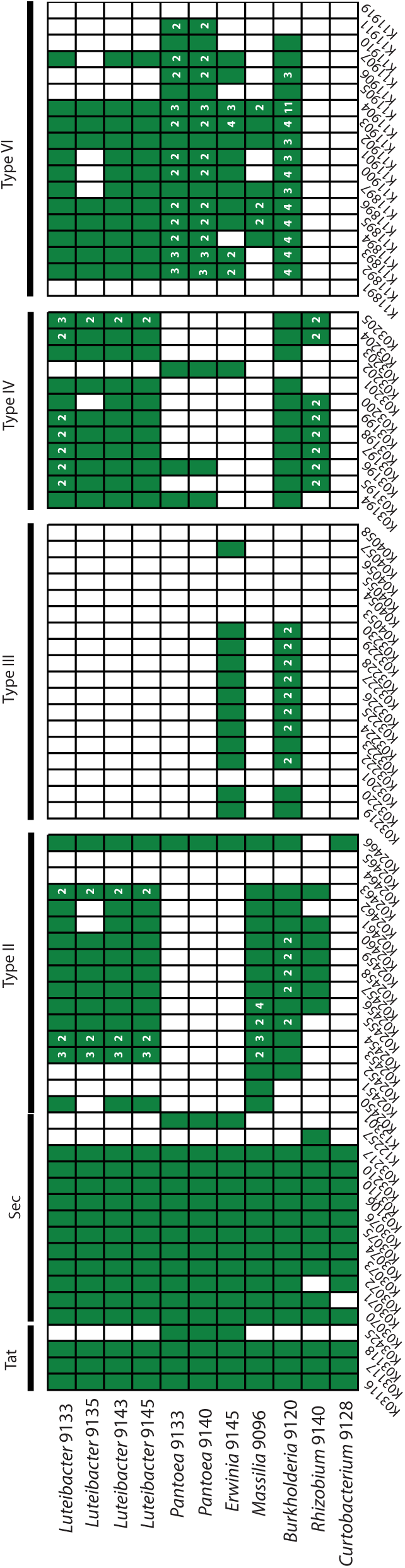
Lack of Conservation Across Facultative Endohyphal Bacteria of Structures Implicated in Interkingdom Interactions. KEGG pathway searches were implemented in IMG to identify bacterial pathways known to be involved in signaling between bacteria and eukaryotes. The 11 focal genomes of this report are listed across the Y axis. Boxes along the X axis indicate KEGG pathway identifiers for constituent genes for each bacteria secretion system with grouping by system denoted at the top. Green boxes indicate that at least one gene within the genome is present and classified according to that specific KEGG identifier. White numbers inside the green boxes denote that more than one gene within that genome is classified according to that KEGG identifier.

The situation is more complicated in regards to “translocation" based systems. Genomes of only two EHB (*Erwinia* sp. 9145 and *Burkholderia* sp. 9120) appear to encode type III secretion systems, with the *Burkholderia* genome likely encoding two separate systems. Type IV systems are found throughout many of these genomes, with *Luteibacter* sp. 9133 and *Rhizobium* sp. 9140 appearing to encode two separate systems. Since the *Luteibacter* genomes each assemble into one contig, it is likely that there are no plasmids present within these strains and therefore that the type IV secretion systems are encoded by the chromosome. In contrast, the genome sequence for the *Rhizobium* strain is split into 7 distinct contigs, which is expected because related strains contain multiple secondary replicons. However, that both type IV systems are present on smaller plasmids suggests that they encode a plasmid transfer system.

The type VI systems present across these genomes are the most complicated to characterize. On one hand, all focal *Luteibacter* strains and the *Erwinia* strain appear to encode one type VI system, whereas the *Pantoea* strains appear to encode two distinct systems on the main chromosome. The *Burkholderia* strain appears to encode four separate systems and 11 different VgrG proteins. Three of these systems are encoded by the main chromosome, whereas one appears to be on a smaller contig (likely a plasmid or mini-chromosome).

## Conclusions

Herein we report nearly complete genome sequences for a diverse suite of bacteria found living inside fungal hyphae. Phylogenetic analyses suggest that these endohyphal bacteria are distinct from previously described EHB, and that the endofungal lifestyle has convergently and independently evolved over short time scales both across diverse bacterial lineages, and in closely related taxa such as *Pantoea* and *Erwinia*. We evaluated the presence/absence of structures known to be involved in interkingdom interactions between bacteria and eukaryotes. Although each strain contains structures that could mediate interactions with fungi, no general mechanism appears to be conserved across strains.

More broadly, these genome sequences provide insights into the ecology of these facultative EHB. Compared to previously characterized genomes from EHB, these sequences do not display dramatic size reductions compared to closely related strains isolated from environmental sources other than fungi. Furthermore, we find that different classes of fungi harbor very similar bacteria. Therefore, these genome sequences are consistent with previous suggestions that these facultative EHB are horizontally transmitted across diverse fungal hosts in nature (Hoffman & Arnold, 2010; see also Arendt *et al*., 2016).

Fungal endophytes, and the Ascomycota that they represent, are largely thought to be hyperdiverse. Our results suggest that diverse bacteria have gained the apparati needed to infect these ecologically and economically important fungi. Given the capacity of EHB to influence fungal phenotypes, these findings illuminate a new dimension of fungal biodiversity.

## Acknowledgments

We thank the National Science Foundation (DEB-1045766 and DEB-0702825 to AEA; IOS-1354219 to DAB, AEA, and Rachel E. Gallery), and the College of Agriculture and Life Sciences at the UA (DAB, KRA, AEA) for financial support. The work conducted by the U.S. Department of Energy Joint Genome Institute, a DOE Office of Science User Facility, is supported by the Office of Science of the U.S. Department of Energy under Contract No. DE-AC02-05CH11231.

## References

Arendt, K.R. (2015) Symbiosis establishment and ecological effects of endohyphal bacteria on foliar fungi. MS Thesis, University of Arizona

Arendt, K. R., Hockett, K. L., Araldi-Brondolo, S. J., Baltrus, D. A. & Arnold, A. E. (2016). Isolation of endohyphal bacteria from foliar Ascomycota and in vitro establishment of their symbiotic associations. Appl Environ Microbiol AEM. 00452–16.

Baltrus, D. A., Dougherty, K., Beckstrom-Sternberg, S. M, Beckstrom-Sternberg, J. S. & Foster, J. T. (2014). Incongruence between multi-locus sequence analysis (MLSA) and whole-genome-based phylogenies: Pseudomonas syringae pathovar pisi as a cautionary tale. Mol Plant Pathol 15, 461–465.

Barre, A. (2004). MolliGen, a database dedicated to the comparative genomics of Mollicutes. Nucleic Acids Research 32, 307D–310.

Bertels, F., Silander, O. K., Pachkov, M., Rainey, P. B. & van Nimwegen, E. (2014). Automated reconstruction of whole-genome phylogenies from short-sequence reads. Mol Biol Evol 31, 1077–1088.

Burstein, D., Amaro, F., Zusman, T., Lifshitz, Z., Cohen, O., Gilbert, J.A., Pupko, T., Shuman, H.A., & Segal, G. (2016). Genomic Analysis of 38 Legionella species identifies large and diverse effector repertoires. Nat. Genet. 48(2): 167–175

Chin, C.-S., Alexander, D. H., Marks, P., Klammer, A. A., Drake, J., Heiner, C., Clum, A., Copeland, A., Huddleston, J. & other authors. (2013). Nonhybrid, finished microbial genome assemblies from long-read SMRT sequencing data. Nat Meth 10, 563–569.

Costa, T. R. D., Felisberto-Rodrigues, C., Meir, A., Prevost, M. S., Redzej, A., Trokter, M. & Waksman, G. (2015). Secretion systems in Gram-negative bacteria: structural and mechanistic insights. Nat Rev Micro 13, 343–359.

Martinez-Cano, D.J., Reyes-Prieto, M., Martinez-Romero, E., Partida-Martinez, L.P., Latorre, A., Moya, A., & Delaye, L. (2015). Evolution of small prokaryotic genomes Front Microbiol 6:742 doi: 10.3389/fmicb.2014.00742

Delepelaire, P. (2004). Type I secretion in gram-negative bacteria. Biochimica et Biophysica Acta 1694, 149–161.

Eid, J., Fehr, A., Gray, J., Luong, K., Lyle, J., Otto, G., Peluso, P., Rank, D., Baybayan, P. & other authors. (2009). Real-time DNA sequencing from single polymerase molecules. Science 323, 133–138.

Ghignone, S., Salvioli, A., Anca, I., Lumini, E., Ortu, G., Petiti, L., Cruveiller, S. E. P., Bianciotto, V., Pietro Piffanelli & other authors. (2011). The genome of the obligate endobacterium of an AM fungus reveals an interphylum network of nutritional interactions. ISME J 6, 136–145.

Grote, J., Thrash, J. C., Huggett, M. J., Landry, Z. C., Carini, P., Giovannoni, S. J. & Rappe, M. S. (2012). Streamlining and Core Genome Conservation among Highly Divergent Members of the SAR11 Clade. MBio 3, e00252–12–e00252–12.

Hoffman, M. T. & Arnold, A. E. (2010). Diverse Bacteria Inhabit Living Hyphae of Phylogenetically Diverse Fungal Endophytes. Appl Environ Microbiol 76, 4063–4075.

Huntemann, M., Ivanova, N. N., Mavromatis, K., Tripp, H. J., Paez-Espino, D., Palaniappan, K., Szeto, E., Pillay, M., Chen, I.-M. A. & other authors. (2015). The standard operating procedure of the DOE-JGI Microbial Genome Annotation Pipeline (MGAP v.4). Standards in Genomic Sciences 1–6. Standards in Genomic Sciences.

Kearse, M., Moir, R., Wilson, A., Stones-Havas, S., Cheung, M., Sturrock, S., Buxton, S., Cooper, A., Markowitz, S. & other authors. (2012). Geneious Basic: An integrated and extendable desktop software platform for the organization and analysis of sequence data. Bioinformatics 28, 1647–1649.

Korotkov, K. V., Sandkvist, M. & Hol, W. G. J. (2012). The type II secretion system:biogenesis, molecular architectureand mechanism. Nat Rev Micro 10, 336–351.

Lackner, G., Moebius, N. & Hertweck, C. (2010). Endofungal bacterium controls its host by an hrp type III secretion system. ISME J 5, 252–261.

Lackner, G., Moebius, N., Partida-Martinez, L. P., Boland, S. & Hertweck, C. (2011). Evolution of an endofungal lifestyle: Deductions from the Burkholderia rhizoxinica genome. BMC Genomics 12, 210.

Lowe, T. M. & Eddy, S. R. (1997). tRNAscan-424 SE: a program for improved detection of transfer RNA genes in genomic sequence. Nucleic Acids Research 25, 955–964.

Markowitz, V. M., Mavromatis, K., Ivanova, N. N., Chen, I.-M. A., Chu, K. & Kyrpides, N. C. (2009). IMG ER: a system for microbial genome annotation expert review and curation. Bioinformatics 25, 2271–2278.

Matz, C., Moreno, A. M., Alhede, M., Manefield, M., Hauser, A. R., Givskov, M. & Kjelleberg, S. (2008). Pseudomonas aeruginosa uses type III secretion system to kill biofilm-associated amoebae. ISME J 2, 843–852.

Naito, M., Morton, J. B. & Pawlowska, T. E. (2015). Minimal genomes of mycoplasmarelated endobacteria are plastic and contain host-derived genes for sustained life within Glomeromycota. Proc Natl Acad Sci USA 112, 7791–7796.

Nazir, R., Hansen, M. A., Sorensen, S. & van Elsas, J. D. (2012). Draft Genome Sequence of the Soil Bacterium Burkholderia terrae Strain BS001, Which Interacts with Fungal Surface Structures. J Bacteriol 194, 4480–4481.

Partida-Martinez, L. P. & Hertweck, C. (2005). Pathogenic fungus harbours endosymbiotic bacteria for toxin production. Nat Cell Biol 437, 884–888.

Pati, A., Ivanova, N. N., Mikhailova, N., Ovchinnikova, G., Hooper, S. D., Lykidis, A. & Kyrpides, N. C. (2010). GenePrimP: a gene prediction improvement pipeline for prokaryotic genomes. Nat Meth 7, 455–457.

Pruesse, E., Quast, C., Knittel, K., Fuchs, B. M., Ludwig, W., Peplies, J. & Glöckner, F. O. (2007). tRNAscan-424 SE: a program for improved detection of transfer RNA genes in genomic sequence. Nucleic Acids Research 35, 7188–7196.

Sachs, J. L., Skophammer, R. G. & Regus, J. U. (2011). Evolutionary transitions in bacterial symbiosis. Proc Nat Acad Sci USA 108 Suppl 2, 10800–10807.

Salvioli, A., Ghignone, S., Novero, M., Navazio, L., Venice, F., Bagnaresi, P. & Bonfante, P. (2015). Symbiosis with an endobacterium increases the fitness of a mycorrhizal fungus, raising its bioenergetic potential ISMEJ 10, 130–144.

Schulz, F. & Horn, M. (2015). SILVA: a comprehensive online resource for quality checked and aligned ribosomal RNA sequence data compatible with ARB. Trends Cell Biol 25, 339–346. Elsevier Ltd.

Torres-Cortés, G., Ghignone, S., Bonfante, P. & Schüßler, A. (2015). Evolutionary transitions in bacterial symbiosis. Proc Nat Acad Sci USA 112, 7785–7790.

Van der Henst, C., Scrignari, T., Maclachlan, C. & Blokesch, M. (2015). An intracellular replication niche for Vibrio cholera in the amoeba Acanthamoeba castellanii ISMEJ 10, 897–910.

van Elsas, J. D. (2014). Burkholderia terrae BS001 migrates proficiently with diverse fungal hosts through soil and provides protection from antifungal agents Front Microbiol 5:598 doi: 10.3389/fmicb.2014.00598

